# Inference of phenotype-relevant transcriptional regulatory networks elucidates cancer type-specific regulatory mechanisms in a pan-cancer study

**DOI:** 10.1101/389734

**Authors:** Amin Emad, Saurabh Sinha

## Abstract

Reconstruction of transcriptional regulatory networks (TRNs) is a powerful approach to unravel the gene expression programs involved in healthy and disease states of a cell. However, these networks are usually reconstructed independent of the phenotypic properties of the samples and therefore cannot identify regulatory mechanisms that are related to a phenotypic outcome of interest. In this study, we developed a new method called InPheRNo to identify ‘phenotype-relevant’ transcriptional regulatory networks. This method is based on a probabilistic graphical model whose conditional probability distributions model the simultaneous effects of multiple transcription factors (TFs) on their target genes as well as the statistical relationship between target gene expression and phenotype. Extensive comparison of InPheRNo with related approaches using primary tumor samples of 18 cancer types from The Cancer Genome Atlas revealed that InPheRNo can accurately reconstruct cancer type-relevant TRNs and identify cancer driver TFs. In addition, survival analysis revealed that the activity level of TFs with many target genes could distinguish patients with good prognosis from those with poor prognosis.

## INTRODUCTION

Gene expression programs are responsible for many biological processes in a cell and extensive efforts have been devoted to elucidating these programs in healthy and disease states. Transcriptional regulatory networks (TRNs) have proven to be a useful framework for describing expression programs. A TRN is a network with transcription factors (TFs) and genes as nodes where a TF-gene edge represents a regulatory effect of the TF on the gene. TRNs are usually constructed from transcriptomic data across many conditions, alone or in combination with other data types. We are especially interested in methods for TRN reconstruction from expression data alone, due to their broad applicability. The majority of such methods are agnostic of any phenotypic annotations of sampled conditions (e.g., case versus control status in disease studies, or drug sensitivity of cell lines in pharmacogenomics studies), looking only to capture correlations between TF and gene expression values in those conditions [1–3]. As a result, many edges in the reconstructed networks may not be relevant to the phenotype being investigated by expression profiling. To take a simple example, consider the two scenarios of gene expression relationship between TF and gene shown in Figure 1A and 1B. In both cases, a linear relationship is evident and is often interpreted as evidence for a TF-gene edge in the TRN. However, it is also apparent that the TF-gene relationship is potentially informative about the phenotypic class in the example of Figure 1B and likely to be irrelevant to the phenotype in the other example (Figure 1A). We believe there is a clear need for methods to reconstruct TRNs that are more focused on phenotype-relevant regulatory interactions (similar to Figure 1B) and can help explain phenotypic variations such as response to different cytotoxic treatments, patient survival, cancer subtypes, etc.

**Figure 1:**
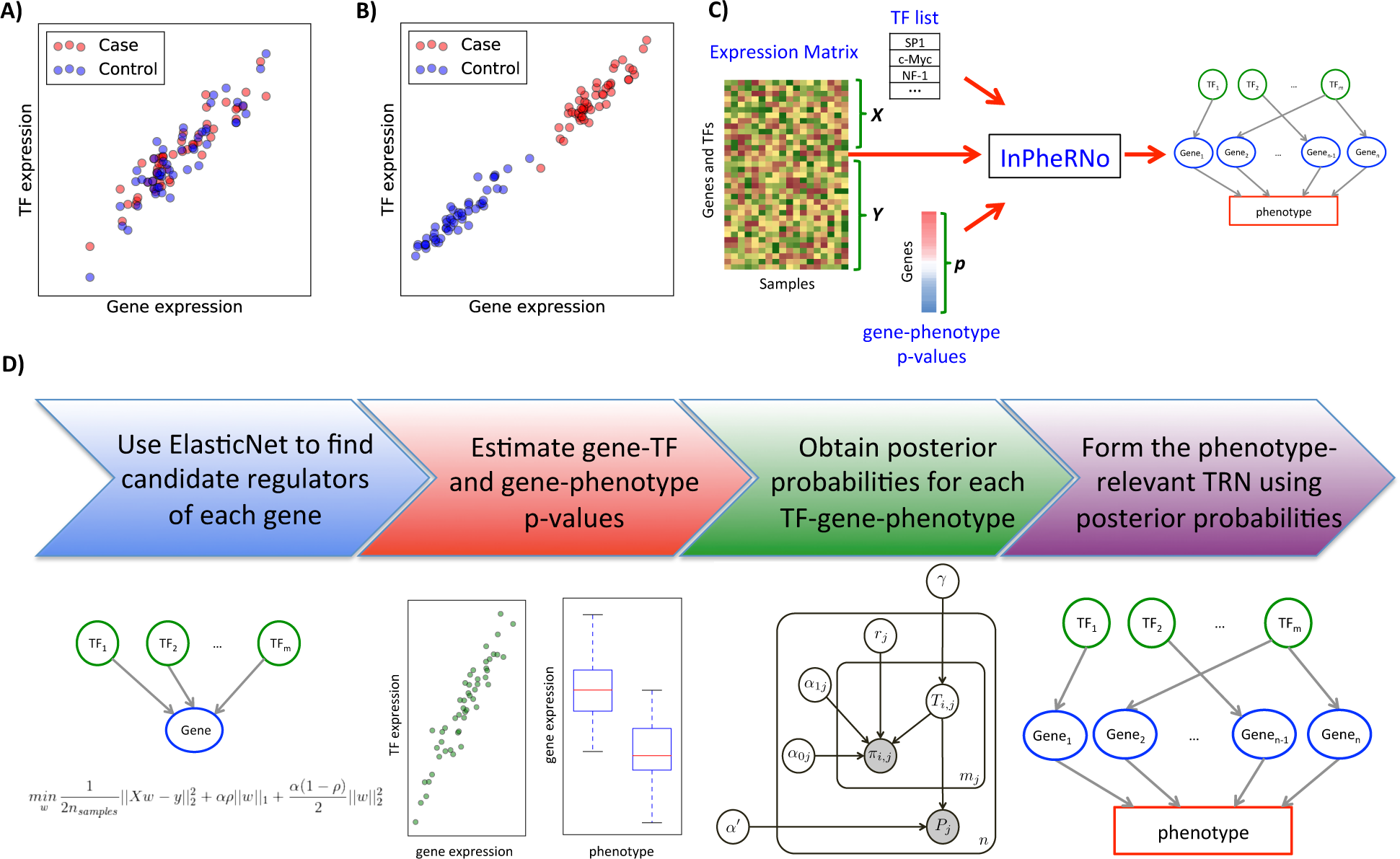
The phenotype-relevant TRN concept and an overview of the InPheRNo framework. (A) The gene-TF expression correlation (across different samples) is independent of the phenotype variation. (B) The gene-TF expression correlation is phenotype-relevant. (C) The inputs and outputs to InPheRNo are shown. The inputs include a matrix of gene expression for all genes (including TF genes), a list of TFs and a vector containing p-value of gene-phenotype associations, denoted as *p.* The list of TFs is used to divide the expression matrix into a matrix **X** of TF expressions and a matrix **Y** of gene expressions. As the output InPheRNo provides a phenotype-relevant TRN. (D) An overview of the InPheRNo pipeline is shown. First, the expression of genes and TFs are used in an Elastic Net algorithm to reduce the number of candidate TFs for each gene. Then, the pseudo p-value of association between TF *i* and gene *j* (denoted by *π_i,j_*) is estimated using an OLS regression model that relates the expression of gene *j* to the expression of m_*j*_ candidate TFs. In addition, the p-values of gene-phenotype associations (denoted by *P_j_*) are assumed to be estimated and provided through *p* for *n* genes. These sets of p-values are used as observed variables in a probabilistic graphical model to learn posterior probabilities for the (TF, gene, phenotype) triplets that a TF regulates a gene to affect the phenotype. These posterior probabilities are used to form the phenotype-relevant TRN.

One approach for including phenotypic information in regulatory network reconstruction is to restrict the analysis to samples representing a particular biological context (e.g. a tissue type [4, 5] or a cancer type [6, 7]). While this approach, henceforth called ‘context-restricted’ TRN reconstruction, may identify important regulatory mechanisms relevant to a context, it does not solve the problem mentioned above - to reconstruct TRNs that explain the variation in a phenotypic outcome. ‘Differential network analysis’ is another approach to relate TRNs to the phenotypic variation. Here, two context-restricted networks are reconstructed based on samples from each of two phenotypic classes, e.g., case versus control, and a differential network is formed by comparing these two networks [8–12]. In focusing on the differential topology of regulatory networks, such methods may fail to identify important phenotype-relevant regulatory edges. For example, Figure 1B illustrates a TF-gene relationship that qualifies as being ‘phenotype-relevant’ and perturbations that abolish it might affect the phenotype; however, such pairs are discarded by the differential network analysis. In addition, these methods cannot be used with continuous-valued phenotypes, and become cumbersome even for categorical phenotypes with more than two categories. A third class of methods is that of ‘context-specific’ network analysis, in which genes associated with phenotype variation are identified, e.g., by differential expression analysis, and then a network is constructed by relating the expression of these genes to the expression of TFs [13–15]. However, one major disadvantage of this approach is that the phenotype-relevance of genes is simply used as a filtering criterion based on arbitrary thresholds and its strength is ignored in TRN reconstruction. Finally, we note that several methods directly evaluate the association between the phenotypic variation and molecular characteristics, including but not limited to gene expression, to identify genes, TFs or miRNAs that can explain the phenotypic variation [16–18]. These methods, while useful for understanding the molecular mechanisms of phenotypic differences, do not directly address the problem of reconstructing phenotype-relevant TRNs. In summary, TRNs are a highly useful and widely popular construct for characterizing gene expression programs underlying phenotypes, yet there is an urgent need for methods that incorporate phenotypic information directly into TRN reconstruction.

We report here a new computational method called InPheRNo (<u>In</u>ference of <u>Phe</u>notype-relevant <u>R</u>egulatory <u>N</u>etw<u>o</u>rks) to reconstruct TRNs that help explain the variation in a phenotype of interest. It models the simultaneous effect of multiple TFs on their targets, as well as the target genes’ association with the phenotype. Its rigorous probabilistic model can be used with categorical or continuous-valued phenotypes, and also provides a confidence score for the identified TF-gene regulatory edges. We applied InPheRNo to data from The Cancer Genome Atlas (TCGA) pertaining to 18 different cancer types, to reconstruct TRNs that differentiate one cancer type from other types of cancer. We also compared them to tissue-specific TRNs reconstructed by analysis of expression data from the GTEx project [19], in order to make the former more specific to the cancer type. The resulting cancer type-relevant TRNs identified regulatory mechanisms involved in the development and progress of each cancer type and discerned previously known as well as novel cancer driver TFs that could be used as potential drug targets. In addition, survival analysis revealed that a gene expression signature formed using these TFs and their target genes can accurately distinguish between patients with poor prognosis and those with good prognosis for the majority of the cancer types. We demonstrated the improved accuracy of InPherNo-derived networks by comparing them to several baseline methods with respect to the above tasks of driver TF discovery and survival prediction. As transcriptomic profiling becomes a standard tool in the study of phenotypic variation among individuals [20], the new tool presented here will help distil the associated high dimensional information into specific regulatory mechanisms underlying that variation.

## RESULTS

### A new probabilistic method for phenotype-relevant transcriptional regulatory network reconstruction

We developed a new computational method called InPheRNo to reconstruct phenotype-relevant transcriptional regulatory networks (TRNs). It analyzes gene expression profiles of a set of samples, along with associated phenotypic scores or labels of those samples, to report TF-gene regulatory relationships relevant to the phenotype. The method is outlined in Figures 1C-1D, and explained in Methods. We touch upon its main steps here. Given the expression of genes and TFs across different samples, first a regression model is used to predict each gene’s expression as a weighted sum of TF expression values. This step uses the Elastic Net regression model [21], which automatically selects a small number of candidate TFs regulating each gene. Next, an ordinary least squares (OLS) regression model is used obtain a pseudo p-value reflecting the importance of each TF to that gene. Note that both of the previous steps use multi-variable regression and model a gene’s expression with a combination of TFs rather than one TF at a time. Separately, a p-value of association between the gene’s expression and the phenotype is obtained using a suitable statistical test. This step allows for different types of phenotypic scores, including categorical labels with two or more values as well as numeric scores, to be incorporated into the method since the gene-phenotype relationship only needs to be encapsulated in a p-value. The two sets of p-values from the above steps – one capturing TF-gene regulatory relationships and the other gene-phenotype associations – are then used as observed variables in a probabilistic graphical model (PGM). The PGM has a latent binary variable for each TF-gene pair, indicating whether the TF regulates the gene so as to affect the phenotype. A Markov chain Monte Carlo (MCMC) algorithm is used to estimate posterior probabilities for these latent variables; these posterior probabilities are then used to obtain a phenotype-relevant TRN (see Methods for more details).

It is worth mentioning that InPheRNo considers the simultaneous effect of multiple TFs on each gene in several steps: first, it utilizes a multivariable Elastic Net model relating the expression of multiple TFs to the expression of the target gene in the TF selection step. Then it obtains a pseudo p-value for each TF-gene pair using a multivariable OLS model, which includes the expression of all selected TFs. Finally, for each gene the PGM models the relationship of observed data to the latent variables representing all selected TFs simultaneously.

### InPheRNo identifies cancer type-relevant TRNs in a pan-cancer study

We applied InPheRNo to the gene expression profiles of 6,357 primary tumor samples corresponding to 18 different cancer types from TCGA, downloaded from the Genomic Data Commons [22], to reconstruct TRNs that differentiate one cancer type from all others. The name, abbreviation and the number of samples of each cancer type used in this study are provided in Table 1. For each cancer type, the phenotype of a sample was defined as a binary variable representing whether the sample is from the cancer of interest or not. The cancer type-relevant TRNs are provided in Supplementary File S1 and the extent of shared regulatory edges between each pair of cancer types is shown in Figure 2A. By and large, the TRNs are noted as being specific to each cancer type (average Jaccard coefficient of shared edges is 0.12), though the pairs (ACC, PCPG), (LGG, GBM), (LUAD, LUSC) and (COAD, READ) exhibit relatively large sharing of edges, partly due to their same tissues of origin. Also noticeable is the high degree of edge-sharing among STAD, COAD, READ and ESCA, all of which are gastro-intestinal cancers.

**Figure 2:**
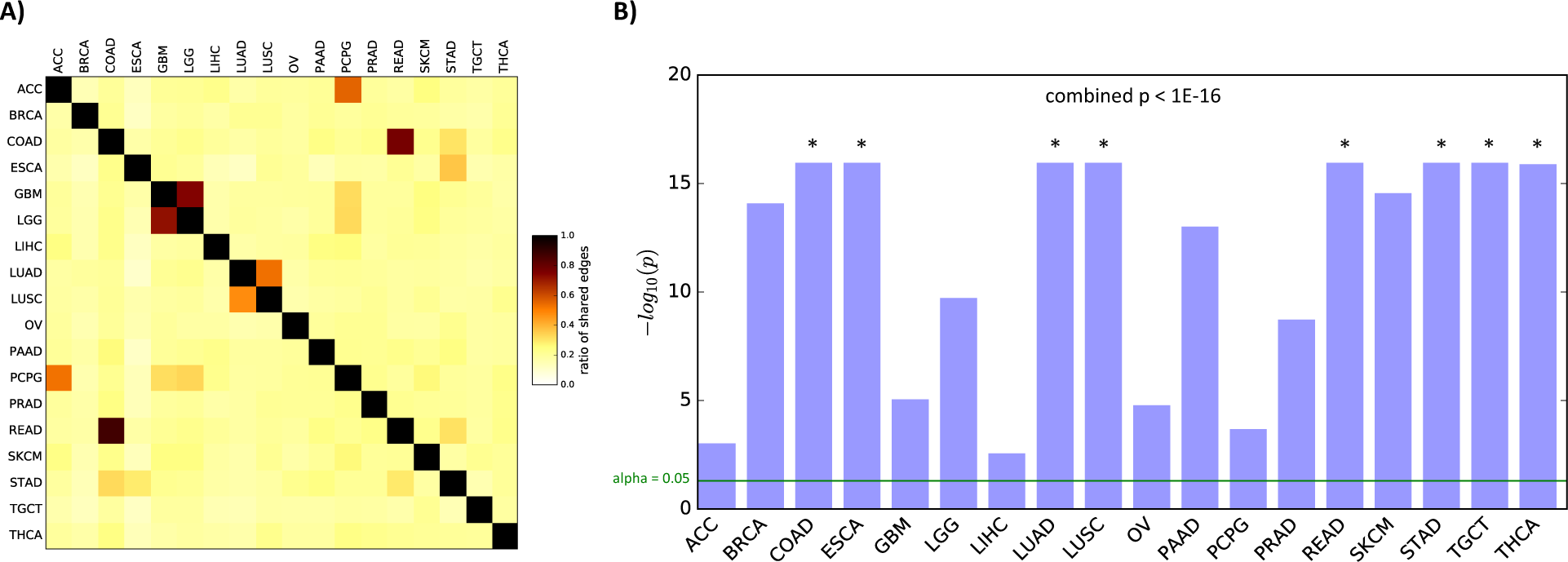
Characteristics of the cancer-relevant regulatory edges identified using TCGA data on 18 cancer types. (A) The heatmap shows the ratio of the shared regulatory edges between a pair of cancers to the total number of edges. More precisely, for any two cancers C_i_ and C_j_, the value in cell (i, j) shows the number of shared regulatory edges divided by the number of regulatory edges in C_i_. (B) The overlap between InPheRNo-identified TRNs for different cancers and global TRNs identified by TREG. The bars represent −log_10_(*p*) of enrichment (hypergeometric test), truncated at 1E-16. The green line shows the threshold alpha = 0.05 and the symbol * is used for cases in which p < 1E-16. The combined p-value is calculated using Fisher‘s method.

**Table 1:** Name, abbreviation and number of samples for each cancer type used in this study.

| Name of the cancer | Abbreviation | Number of Samples |
| --- | --- | --- |
| Adrenocortical carcinoma | ACC | 79 |
| Brain Lower Grade Glioma | LGG | 511 |
| Breast invasive carcinoma | BRCA | 1091 |
| Colon adenocarcinoma | COAD | 456 |
| Esophageal carcinoma | ESCA | 161 |
| Glioblastoma multiforme | GBM | 154 |
| Liver hepatocellular carcinoma | LIHC | 371 |
| Lung adenocarcinoma | LUAD | 513 |
| Lung squamous cell carcinoma | LUSC | 501 |
| Ovarian serous cystadenocarcinoma | OV | 374 |
| Pancreatic adenocarcinoma | PAAD | 177 |
| Pheochromocytoma and Paraganglioma | PCPG | 178 |
| Prostate adenocarcinoma | PRAD | 495 |
| Rectum adenocarcinoma | READ | 166 |
| Skin Cutaneous Melanoma | SKCM | 103 |
| Stomach adenocarcinoma | STAD | 375 |
| Testicular Germ Cell Tumors | TGCT | 150 |
| Thyroid carcinoma | THCA | 502 |

Due to differences in tissues of origin of the studied cancer types, some of the regulatory mechanisms identified as differentiating one cancer from others may reflect these tissue differences and not the cancers themselves. To address this and better characterize cancer type-specific mechanisms, we additionally applied InPheRNo to gene expression data profiles of 4,388 normal tissue samples in the Genotype-Tissue Expression Project (GTEx) data portal [19], corresponding to the 18 cancer types above (Supplementary Table S1 in Supplementary File S2). The identified tissue-relevant TRNs (Supplementary File S3) should enable us to distinguish between regulatory mechanisms in a normal tissue from regulatory mechanisms involved in a cancer originating from that tissue, a direction we pursue later.

As a preliminary assessment of their accuracy, we sought to determine whether the identified cancer type-relevant TRN edges are enriched in independently identified TF-gene relationships. Although the TRNs derived above are meant to be phenotype-relevant, they reflect regulatory relationships and are thus expected to be enriched in globally characterized regulatory edges, albeit to different degrees depending on the specific cancer. We therefore used global TRNs reconstructed from ChIP-seq profiles of 166 TFs and matched gene expression data in 43 different cell lines from the ENCODE project, using the TREG method [23] (see Methods for details). Figure 2B and supplementary Figure S1 (in Supplementary File S2) show the extent to which the cancer type-relevant TRN edges identified using InPheRNo are enriched for global TRN edges. As expected, we observed significant enrichments for every cancer type, but to different degrees. Similarly, for all tissues expect one, tissue-relevant regulatory edges obtained by applying InPheRNo on GTEx data are enriched in global regulatory edges (Supplementary Figure S1 in Supplementary File S2).

We noted a significant correlation between different cancer types and their corresponding normal tissues in terms of their enrichment for global TRN edges (Spearman’s rank correlation = 0.63, p = 4.8E-3). This is in line with our expectation that some of the regulatory mechanisms identified from the TCGA data reflect the differences in regulatory mechanisms of the tissues of origin. To correct for this confounding effect, for each cancer we removed all the edges that were also present in the TRN identified for its corresponding normal tissue (Supplementary File S4). Depending on the cancer type, this procedure removed 7.0% (for READ) to 10.3% (for LUSC) of the identified edges (Supplementary Figure S2 in Supplementary File S2). The analyses reported in the rest of the manuscript correspond to these tissue-corrected results.

### InPheRNo identifies breast cancer-relevant ‘driver’ transcription factors

It is challenging to assess the accuracy and cancer-relevance of predicted TF-gene relationships on a global scale. However, TFs with many target genes in our cancer-relevant TRNs are expected to play important roles in cancer origin and progression, and existing databases of cancer drivers may therefore help us evaluate the TRNs. Accordingly, we examined the concordance between key TFs identified in the cancer type-relevant TRNs above and known driver TFs for that cancer as catalogued in the DriverDBv2 [24] and IntOGen [25] databases. We first focused on breast cancer (BRCA) given the relatively extensive knowledge of driver genes for it. We examined the BRCA-relevant TRN reconstructed using InPheRNo (Figure 3A) and identified 15 TFs with most targets (Table 2) in this network. This set included six BRCA driver TFs (RUNX1, GATA3, MYB, FOXA1, ZBTB41, PRRX1) according to DriverDBv2 [24] (p = 1.2E-4, hypergeometric test) and four according to IntOGen [25] (p = 2.3E-4). To assess if the InPheRNo TRNs exhibit an improved ability to reveal driver TFs, we repeated the above evaluations with results from five alternative approaches (see Methods for details of each approach), as outlined below.

**Table 2:**
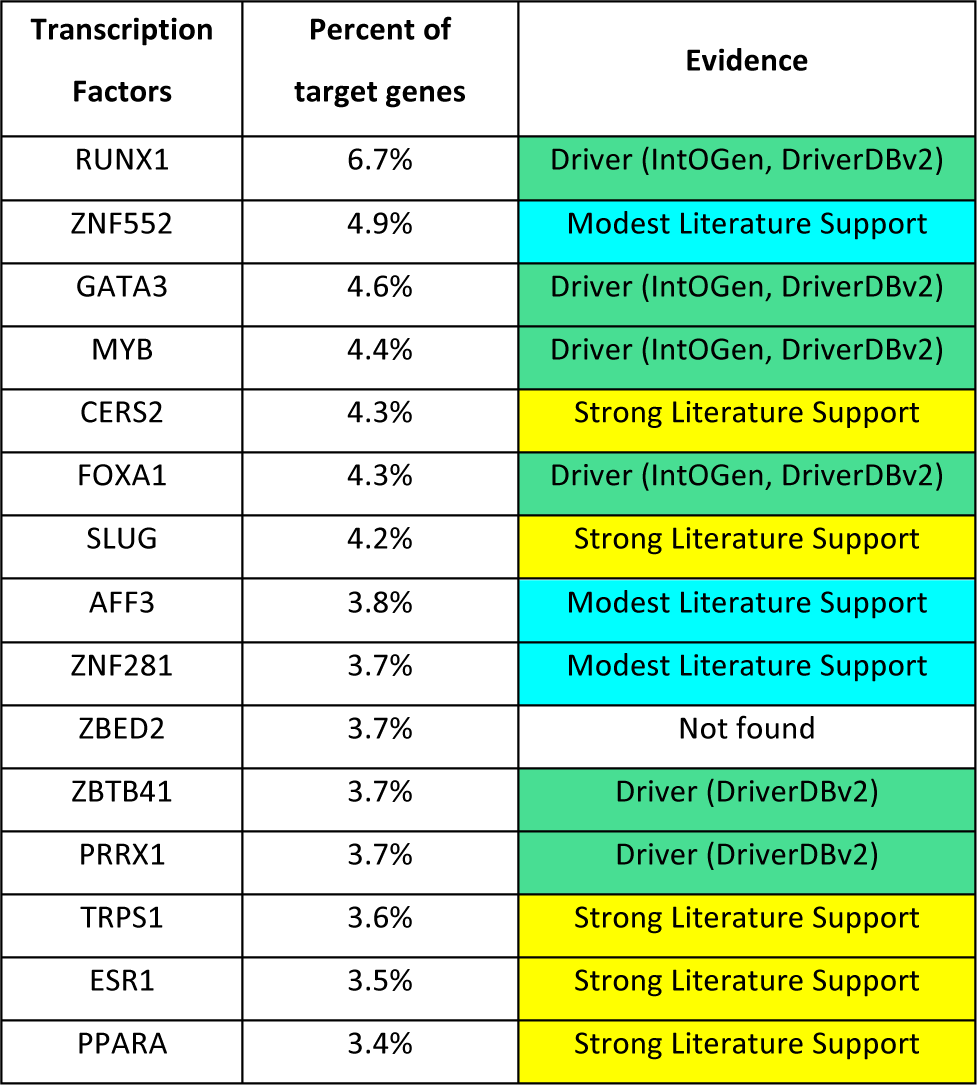
Top 15 TFs identified using InPheRNo and the evidence for their role in Breast Cancer. The TFs are ranked based on the number of their cancer-relevant target genes. The second column shows the percent of the considered genes that each TF regulates.

| Transcription Factors | Percent of target genes | Evidence |
| --- | --- | --- |
| RUNX1 | 6.7% | Driver (IntOGen, DriverDBv2) |
| ZNF552 | 4.9% | Modest Literature Support |
| GATA3 | 4.6% | Driver (IntOGen, DriverDBv2) |
| MYB | 4.4% | Driver (IntOGen, DriverDBv2) |
| CERS2 | 4.3% | Strong Literature Support |
| FOXA1 | 4.3% | Driver (IntOGen, DriverDBv2) |
| SLUG | 4.2% | Strong Literature Support |
| AFF3 | 3.8% | Modest Literature Support |
| ZNF281 | 3.7% | Modest Literature Support |
| ZBED2 | 3.7% | Not found |
| ZBTB41 | 3.7% | Driver (DriverDBv2) |
| PRRX1 | 3.7% | Driver (DriverDBv2) |
| TRPS1 | 3.6% | Strong Literature Support |
| ESR1 | 3.5% | Strong Literature Support |
| PPARA | 3.4% | Strong Literature Support |

**Figure 3:**
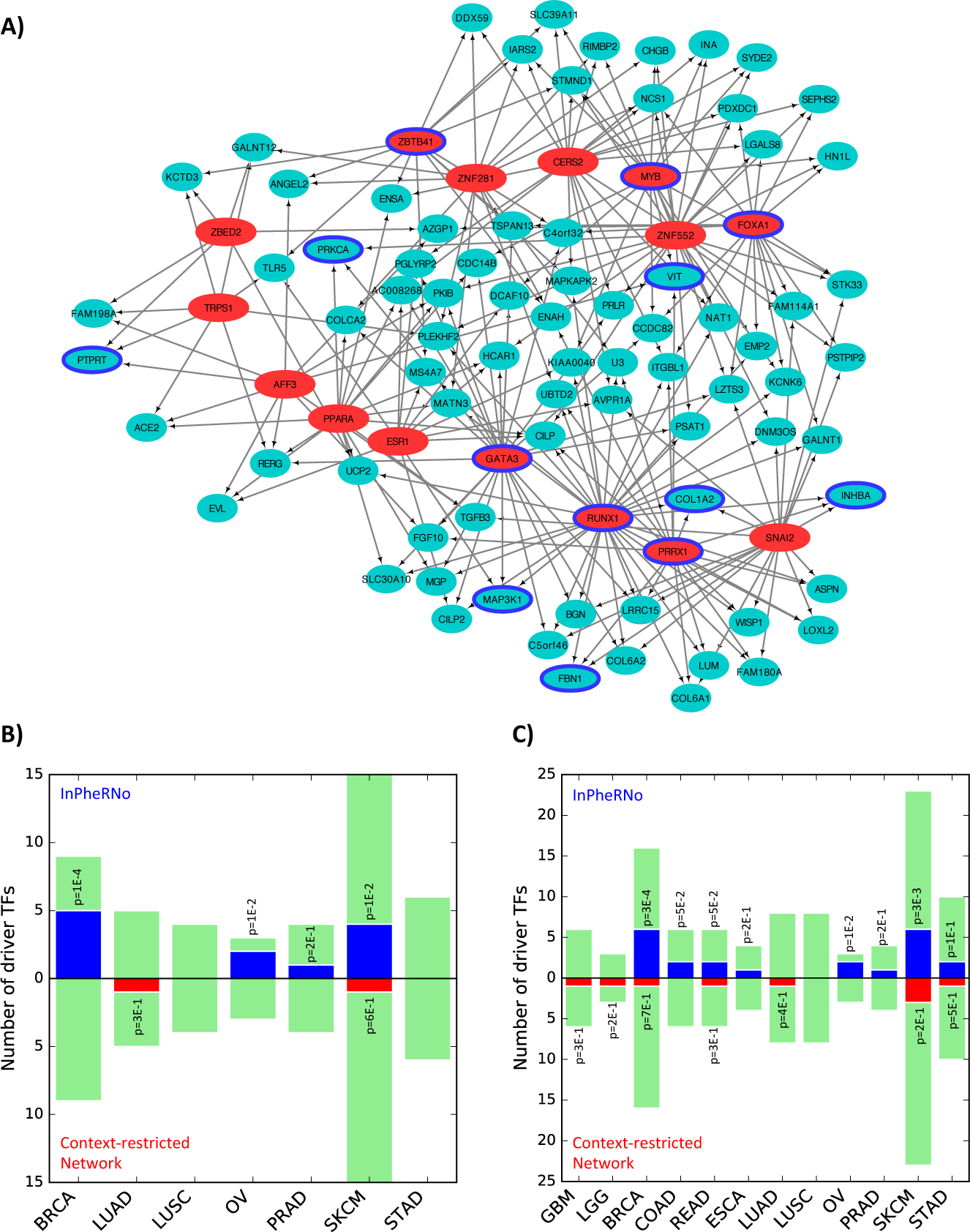
Driver TFs in cancer type-relevant TRNs reconstructed by InPheRNo. (A) A subnetwork of the BRCA-relevant TRN. The depicted subnetwork consists of the 15 TFs (red) with the most target genes, as well as genes (green) that are regulated by at least three of these TFs. Genes or TFs with a blue border represent BRCA drivers according to IntOGen and DriverDBv2. (B-C) Cancer-specificity of InPheRNo in identifying driver TFs (using IntOGen) compared to the context-restricted network analysis. For each cancer type, 100 TFs with the most number of identified target genes are selected and are compared with the set of driver TFs of that cancer that are drivers of at most *n_s_* other cancers. Color green shows the total number of cancer-specific driver TFs in the IntOGen database, color blue corresponds to number of cancer-specific driver TFs identified by InPheRNo and red represents driver TFs identified using context-restricted network analysis. Only cancers that had more than one known cancer-specific driver TF are used for the analysis. The p-values are calculated using a hypergeometric test. (B) Results corresponding to *n_s_ =* 2. (C) Results corresponding to *n_s_ =* 3.

In the first baseline, we constructed a ‘context-restricted’ TRN using only breast cancer samples, mimicking similar approaches in the literature [4–7]. We modeled each gene’s expression in terms of the expression values of all TFs, via multivariable regression. We adopted the Elastic Net algorithm for this purpose, exactly as in the first step of InPheRNo, obtaining a small number of TFs regulating each gene (see Methods), and ranked TFs by the number of target genes. The top 15 TFs identified using this approach included no BRCA-driver TF according to either of the two databases. In the second baseline, we used ‘differential network analysis’ [9] to identify edges that are present in the TRN reconstructed using BRCA samples and not present in the TRN reconstructed using samples of other cancers pooled together (see Methods). (TRN reconstruction relied on the Elastic Net algorithm, exactly as in the first baseline.) The set of 15 TFs with the most number of target genes using this approach contained only one known BRCA-driver TF according to DriverDBv2 and none according to IntOGen. The third baseline was a ‘context-specific’ TRN [13–15] reconstructed by relating the expression of differentially expressed genes to the expression of TFs (see Methods). The set of top 15 TFs identified using this approach did not include any BRCA-driver TFs according to any of the two databases. The fourth method compared involved identifying TFs whose expression had the most significant difference between samples of the breast cancer compared to samples of other cancers (Welch‘s t-test). (That is, no TRN reconstruction was performed.) The set of 15 TFs identified using this approach did not contain any driver TFs according to IntOGen or DriverDBv2. Finally, in the fifth baseline, we used an approach based on Fisher’s method to combine the p-value of the association between a gene’s expression and the phenotype with the p-value of Pearson’s correlation between expression of that gene and the expression of a TF (see Methods for details). This method, which can be considered a simplified version of InPheRNo, has the benefit of reconstructing phenotype-relevant co-expression networks efficiently, but does not allow us to simultaneously model the effect of multiple TFs on each gene. In spite of this shortcoming, this method, henceforth called ‘simplified-InPheRNo’, outperformed all other methods except for InPheRNo in identifying BRCA driver TFs: the list of 15 TFs with the most number of target genes included 4 driver genes according to either database. The top TFs identified using these different methods are provided in Supplementary Table S2.

We noted above that six of the 15 key TFs of the BRCA-specific TRN determined by InPheRNo are known driver TFs. We mined the literature and found strong evidence for the role of five additional TFs (from the remaining nine) in BRCA; see Table 2. For instance, ESR1 encodes estrogen receptor alpha and its role in the development, progress and drug resistance of breast cancer is well documented [26–28]. CERS2 is a ceramide synthase and suppresses breast tumor cell invasion and enhances chemosensitivity of breast cancer cells [29, 30]. In addition, the low expression of this gene is associated with poor prognosis in breast cancer [30]. SLUG is a TF involved in epithelial to mesenchymal transition (EMT) and is known to promote breast cancer progression and invasion [31–33]. We recently showed that this TF (along with FOXA1, another TF identified by InPheRNo, Table 2) is a biomarker of metastatic subtypes of breast cancer [34]. TRPS1 is a transcription repressor of GATA-regulated genes, which promotes EMT in breast cancer and its expression is associated with clinical outcome in this cancer [35, 36]. The activation of PPARA has been shown to promote proliferation in human breast cancer and its genetic polymorphism has been linked to an increase in the odds of postmenopausal breast cancer [37, 38]. In addition to the above five, three other TFs among the top 15 identified by InPheRNo have modest literature support for a role in BRCA development: AFF3 is a nuclear transcriptional activator, which is abnormally expressed in some cases of breast cancer and has been suggested as a proto-oncogene [39, 40]. ZNF281 is a transcriptional repressor involved in EMT that is upregulated in colon and breast cancer and has been suggested to promote these cancers [41, 42]. In addition, ZNF552 has been suggested as a regulator of genetic risk of breast cancer and its regulons have shown to be enriched in genes associated with risk loci identified using a combination of GWAS and eQTL analysis [43]. Taken together, these results suggest that InPheRNo can accurately identify regulatory mechanisms (in this case, major TFs) involved in breast cancer.

### Driver transcription factors identified by InPheRNo are specific to respective cancer types

We next asked if the key TFs (those with most target genes) in InPheRNo-derived TRN_s_ are specific to their respective cancer types, as this is an important criterion for phenotype-relevant TRN reconstruction. We obtained a list of driver TFs for each cancer from IntOGen, and retained only those known drivers that were not annotated as drivers for more than *n_s_* = 2 other cancer types. We then compared these cancer type-specific drivers to the top 100 (out of 1544) TFs identified for that cancer using InPheRNo (Supplementary Table S2). Of the seven cancers types that had more than one known driver TF specific to them, three cancers (BRCA, OV, and SKCM) showed a significant (alpha = 0.05) enrichment between InPheRNo-identified TFs and known cancer type-specific drivers, with an overall combined p-value (Fisher’s method) of p = 2.5E-4 (Figure 3B). However, repeating the above procedure with key TFs identified by context-restricted network analysis, differential network analysis, context-specific network analysis, or based on differential expression did not yield significant enrichment for cancer type-specific drivers in any of these seven cases (Figure 3B and Supplementary Figure S3 in Supplementary File S2). Key TFs of TRNs determined by simplified-InPheRNo were significantly enriched for known drivers in two cases (Supplementary Figure S3 in Supplementary File S2).

Similar observations were made when using a slightly relaxed definition of a cancer type-specific driver TF: as a known driver of one cancer type that is not a known driver for more than *n_s_* = 3 other cancer types (Figure 3C, Supplementary Figure S4 in Supplementary File S2). For the 12 cancer types where two or more such cancer type-specific drivers are known, InPheRNo-identified key TFs showed the highest enrichment for those drivers (combined p = 6.2E-5) compared to simplified-InPheRNo (p = 6.6E-4), top differentially expressed TFs (p = 0.62), differential network analysis (p = 0.64), context-restricted analysis (p =0.92), and context-specific analysis (p=0.99). While the above analyses were performed using driver TF annotations from IntOGen, similar analysis using driver genes in DriverDBv2 also confirmed the conclusion that InPheRNo has a high specificity in identifying regulatory mechanisms involved in each cancer, especially when compared to alternative approaches (Supplementary Figures S5-S7 in Supplementary File S2). We believe this arises from the explicit and quantitative incorporation of cancer type-specificity of target genes into its statistical model.

### Gene expression signatures based on InPheRNo TRNs are predictive of patient survival

Gene expression signature analysis is a widely used approach in analyzing and subtyping cancer samples, with great potential for improving prognosis and treatment [44, 45]. We hypothesized that since InPheRNo identifies cancer type-relevant regulatory mechanisms, the resulting TRNs can be used to form gene expression signatures that are more predictive of patient survival than signatures formed using differential expression analysis, one of the most widely used approaches for forming gene expression signatures [45]. It has been previously suggested that the activity of a TF is better reflected in the activity of its targets than its own expression [16]. Therefore, we formed a gene expression signature for each TF reflecting the expression of the TF as well as the activity levels of its targets in the InPherRNo-derived TRN, while taking into consideration the strength and mode of regulation for each gene (see Methods for details). For each cancer type, we then used signatures of the five key TFs with the largest number of target genes in the corresponding TRN, and clustered patient tumor samples into two groups (hierarchical clustering) based on the resulting signatures. We used Kaplan-Meier survival analysis to determine whether these two clusters show distinct survival behavior, limiting our analysis to cancers with more than 150 samples and ten incidents of death. Out of the thirteen cancers satisfying these conditions, the expression signatures classified samples into clusters of distinct survival (log-rank test, alpha = 0.05) for seven cancers (Figure 4A), with a combined p-value of p = 1.0E-9 (Fisher’s method).

**Figure 4:**
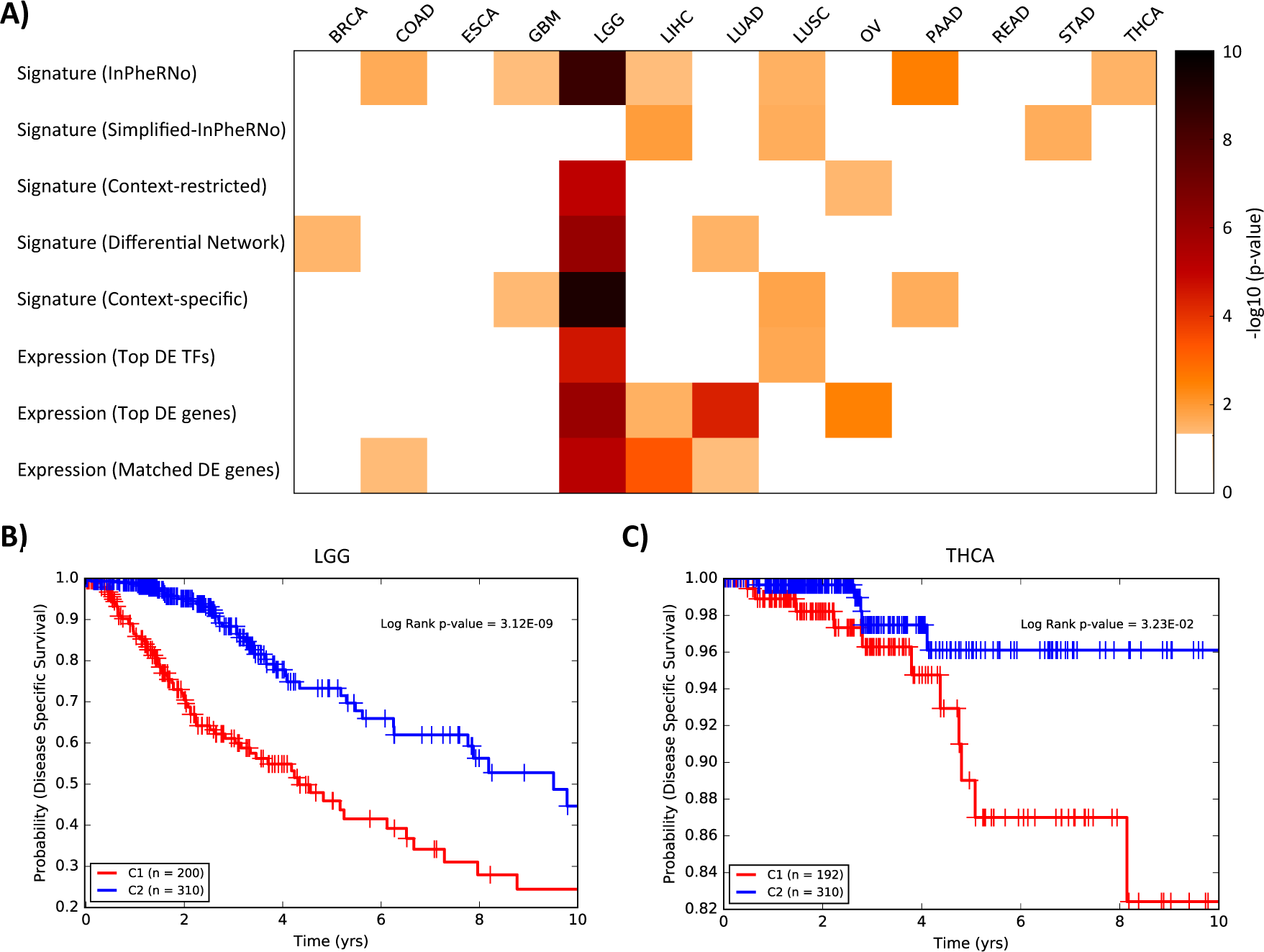
Survival analysis for samples of different cancers clustered using different approaches. (A) The heatmap shows the performance of different approaches used for clustering of samples. Samples of each cancer are clustered into two groups and each cell in the heatmap represents −log_10_(*p*) (obtained using a log-rank test) of the significance of the difference between survival probabilities of the two clusters. For clarity, cases in which the p-value was larger than 0.05 are shown as white. (B-C) Kaplan-Meier analysis for two clusters obtained by the gene expression signature formed by the top 5 TFs and their target genes, as identified by InPheRNo for LGG (B) and THCA (C) cancer types.

We repeated the above survival analysis using gene expression signatures created from TRNs reconstructed by context-restricted analysis, differential network analysis, context-specific analysis and simplified-InPheRNo, which resulted in one to at most four significant cases (Figure 4A, Supplementary Table S3). Similarly, clustering based on top five most significantly differentially expressed genes or TFs resulted in four and two significant cases, respectively. The results did not improve when we used the same number of differentially expressed genes as was used in forming InPheRNo’s gene signature, yielding only four significant cases. These results show that taking into account the phenotype-relevant regulatory mechanisms identified by InPheRNo in developing gene expression signatures can improve the performance of gene signature analysis and prediction of survival.

Given the observation that the gene expression signature formed using the InPheRNo-identified TRN for Lower Grade Glioma (LGG) can accurately predict patients’ prognosis (Figure 4B), we sought to determine the functional characteristics of these genes. To this end, we performed gene ontology (GO) enrichment analysis using KnowEnG analytical platform [46] for each of the five TFs and their targets, one TF at a time. Overall, 49 GO terms with size larger or equal to ten were enriched (Fisher’s exact test, Benjamini-Hochberg corrected false discovery rate p* < 0.05) for these gene sets (Figure 5 and Table S4).

**Figure 5:**
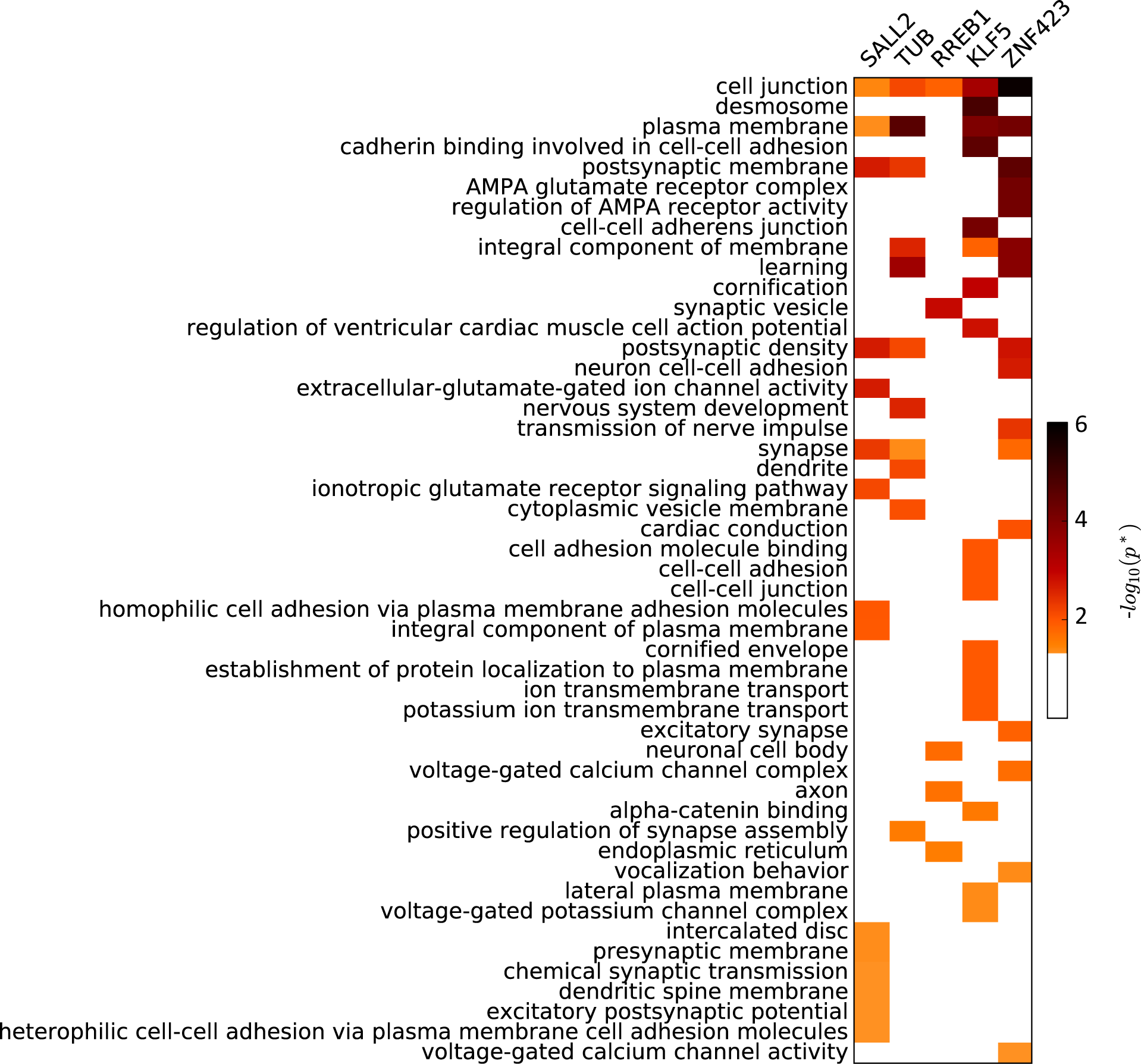
Functional annotation of top 5 TFs and their targets identified using InPheRNo for LGG. The heatmap shows the Benjamini-Hochberg corrected GO enrichment false discovery rates (FDR). For clarity, cases in which the FDR was larger than 0.05 are shown as white. The GO terms are sorted based on the smallest FDR in any of the five gene sets.

Out of these GO terms, 21 were related to the nervous system, neurotransmission, and neurogenesis. On the other hand, 12 terms were related to cell junction, which plays an important role in the invasion-metastasis cascade in various cancers including gliomas [47, 48]. These results support our expectation that both regulatory mechanisms specific to nervous system as well as more general cancer-related mechanisms are involved in the development and progress of LGG.

## DISCUSSION

Transcriptional regulatory networks (TRNs) provide an important and popular framework for better understanding a cell’s regulatory mechanisms in response to or leading to phenotypic conditions. However, TRN reconstruction methods today do not incorporate phenotypic information adequately or at all. As such, the reconstructed networks may be limited in pinpointing regulatory mechanisms most related to a phenotype under investigation, and often necessitate a follow-up step that filters for phenotype-relevance. For example, a recent study of gene expression changes underlying Huntington’s disease (HD) [49] reconstructed a TRN specific to the mouse striatum and then short-listed TFs whose predicted targets were enriched in genes differentially expressed in HD mouse models. In another study, gene expression profiles of TFs and putative target genes were used to reconstruct a context-restricted TRN for breast cancer (using only breast cancer samples), and then a list of breast cancer-relevant TFs (called ‘risk-TFs’) whose regulons were enriched in risk-loci were short-listed [43]. In this study, GWAS and eQTL analyses were used to define risk loci and relate them to the regulon of each TF. Such previous attempts to augment TRN reconstruction with phenotypic data motivated us to develop a systematic approach to incorporate information about the phenotype directly into TRN reconstruction.

In this study, we developed InPheRNo to reconstruct phenotype-relevant TRNs and utilized it to identify regulatory interactions that differentiate one cancer type from others while correcting for the confounding effect of tissues of origin. InPheRNo is based on a carefully designed PGM, which is key to combining TF-gene expression correlations with gene-phenotype associations. The method is broadly applicable since it learns regulatory relationships from expression data alone and does not impose any restriction on the type of phenotype under investigation - the phenotype may be binary, categorical or even continuous-valued, and any appropriate statistical method for testing its association with a gene’s expression may be used in InPheRNo. Unlike several other methods that rely on the regulatory relationship of one TF-gene pair at a time, InPheRNo considers the effect of multiple TFs on each gene in the reconstruction procedure, at the time of selecting candidate TFs as well as in training the PGM. Finally, using posterior probabilities obtained from the PGM, InPheRNo provides a score representing the confidence for the identified phenotype-relevant regulatory edges. Our extensive analyses using a pan-cancer dataset from TCGA showed the advantages of this novel framework compared to other related (yet different) approaches. Our results showed that the TFs with many cancer type-relevant targets are potential cancer driver TFs and may suggest novel drug targets or provide new insights regarding the development and progress of cancer. Our results also suggest a powerful approach for subtyping of cancer patients using gene expression signatures: while most approaches developed for this task do not take into account the regulatory interactions among genes, our survival analysis suggests that cancer-type relevant TRNs can improve the predicting power of gene expression signatures.

In spite of the success of the InPheRNo-based gene signatures in differentiating between patients with poor and good prognosis for the majority of cancer types, in some cases, e.g., BRCA, this method did not result in groups with significantly different survival probability, despite the existence of BRCA-driver TFs in the signature. This lack of success may partially be due to the fact that we clustered samples of each cancer type into two clusters, while these cancer types may include more than two subtypes, as is the case in BRCA [50]. However, since in most cancer types a definite number for the cancer subtypes is not yet established, we preferred to keep the number of clusters equal to two. A more in-depth analysis of subtype discovery and survival analysis using InPheRNo-derived TRNs is left for future work.

Another future direction for improving InPheRNo is to simultaneously include additional types of regulatory evidence, especially those representing ‘cis’ mechanisms such as TF motifs and chromatin state changes, in the TRN reconstruction procedure. This is an important direction, especially since many efforts are under way to generate large datasets containing matching transcriptomic, genomic, epigenomic and phenotypic profiles of many patients. One way to achieve this goal might be to include different regulatory evidence as new observed variables in the PGM used in InPheRNo. However, further investigations are necessary to model the corresponding PGM.

## METHODS

### Inference of Phenotype-relevant Regulatory Networks (InPheRNo)

InPheRNo (Figures 1C–1D) is a new computational method for reconstructing phenotype-relevant TRNs. At its core, InPheRNo utilizes a carefully designed PGM to systematically combine the information on the significance of gene-phenotype associations with the information on the significance of gene-TF associations to obtain a phenotype-relevant TRN. In addition, InPheRNo takes into account the simultaneous effect of multiple TFs on each gene.

As input, InPheRNo accepts a matrix of gene and TF expression data (gene and TFs x samples), a list of TFs and a vector ***p*** that records the p-value of association between the expression of each gene and the phenotypic variation across samples (obtained using a suitable statistical test depending on the type of phenotype), as depicted in Figure 1C. We assume that the expression matrix is properly normalized in advance (described below). Using the list of TFs, the gene expression matrix is divided into a matrix ***X*** of TF expression data (TFs x samples) and a matrix ***Y*** of gene expression data (genes x samples).

In order to obtain a measure of significance for the association between each gene-TF pair, while considering the influence of other TFs on the gene of interest, we used a two-step procedure. First, we used Elastic Net, a linear multivariable regression algorithm that imposes sparsity using regularization, to identify a small set of *m_j_* candidate TFs for each gene *j* (*j* = 1, 2, …, *n*). In this model, the TF expression matrix ***X*** is the feature matrix and the expression profile **y**_*j*_ of each gene is the response vector. The value of *m_j_* is determined by the Elastic Net’s hyperparameters, but the user may select an upper bound *m*_max_ on this value to reduce the running time and impose the prior knowledge that only a few TFs typically regulate each gene. (We used *m*_max_= 15 for our analyses.) Note that the idea of imposing an upper limit on the number of regulators of a gene has been previously used in the literature [51–53].

Next, for each gene *j* we formed a matrix ***X**_j_* representing the expression of the *m_j_* selected TFs across different samples. Then, we used ***X**_j_* as the feature matrix in a multivariable OLS regression model to relate the expression of the identified TFs to the expression of the gene **y**_*j*_ (the response vector) and calculated a pseudo p-value π_*i,j*_ (using a two-sided t-test), reflecting the conditional effect of the TF *i*(*i* = 1, 2,…, *m_j_*) on gene *j.* Using the OLS regression model is a necessary step, since currently no statistical method exists to directly calculate the p-value of feature-response associations in an Elastic
Net model. It is important to note that *π_i,j_* is only a ‘true’ p-value for the second step of this procedure, but does not satisfy all the characteristics of a p-value for the two-step procedure (see Supplementary File S2 for simulation results). More precisely, under the Null hypothesis that TF *i* is not associated with gene *j,* the distribution of *π_i,j_* is not uniform (a characteristic of a true p-value), but instead is biased towards small values (see Supplementary Figures S10-S12 in Supplementary File S2). The reason for this bias is that in the first step, Elastic Net selects TFs whose expression are associated with the expression of gene *j* and the second step is thus likely to assign a small p-value to them. This is an important consideration, since it affects how we model the conditional distributions of *π*_*i,j*_s in the PGM described below.

The two sets of p-values - one capturing TF-gene regulatory relationships (denoted as *π_i,j_*) and the other gene-phenotype associations (denoted as *P_j_* and provided in vector ***p**)* - are used as observed variables in a PGM (Supplementary Figure S8 in Supplementary File S2) that has binary latent variables *T_i,j_* reflecting the role that each putative TF-gene interaction plays in phenotype variation. More precisely, T_*i,j*_ = 1 implies that TF *i* regulates gene *j* so as to afect the phenotype, and *T_i,j_* = 0 indicates its logical complement. We modeled the prior distribution of this random variable as *T_i,j_ ~ Bernoulli*(*γ*). The posterior probabilities of *T_i,j_*s obtained from this PGM can be used to form the phenotype-relevant TRN (as described below).

As depicted in Figure 1D and Supplementary Figure S8 (in Supplementary File S2), InPheRNo uses a causal directed acyclic graph (DAG) to model the relationship between the latent variables and the observed variables. The topology of this DAG represents the idea that the value of *T_i,j_* has a causal effect on the distributions of observed variables *P_j_*s and *π_i,j_*. Since each *P_j_* represents a ‘true’ p-value, it follows a uniform distribution under the Null hypothesis that ‘expression of gene *j* is not associated with the phenotypic variation*',* which is the scenario where gene *j* does not mediate the influence of any of its putative regulators on the phenotype. In other words, if *T*_1*,j*_=*T_2,j_*= … = *T_m,j,_j*= 0, then *P_j_ ~ Unif*(0,1). On the other hand, if any of the *T_i,j_*s is equal to 1, the definition of T_*i,j*_ impliesthat gene *j* is associated with the phenotype (the alternative hypothesis). Following the approach in Hanson et al. [54] who successfully used a Beta distribution to model the distribution of p-values when they are biased towards small values, we used a *Beta*(*α,β*) distribution to model the distribution of these variables under the alternative hypothesis. By fixing *β*= 1 and limiting the value of *α* in the range 0 < *α* ≤ 1, we can obtain a wide range of distributions with different degrees of bias towards small values with the smallest bias when *α* = 1 (equivalent to a uniform distribution) and an increasing degree of bias as *α* approaches 0 (see Supplementary Figure S9 in Supplementary File S2). Thus, the conditional distribution of *P_j_* given the value of its parent nodes in the DAG can be modeled as

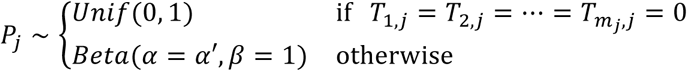

where *α′*, 0 <*α ′*≤ 1, is a parameter controlling the degree of bias of the Beta distribution towards small values. In our analyses, we estimated *α′* by fitting a mixture of a uniform and a Beta distribution to the histogram of *P_j_*s for all genes, prior to training the PGM. Note that modeling the conditional distribution of each *P_j_* with respect to *T_i,j_*s is yet another technique that we used to capture the influence of multiple TFs on the value of observed variables, and hence on the phenotype-relevant regulation of the genes.

As mentioned earlier the pseudo p-values *π_i,j_*s obtained using the two-step procedure are biased towards small values even when TF *i* is not a regulator of gene *j.* As a result, similar to the case with *P_j_*s, we can use two distributions *Beta*(*α*=*α*_1*j*_,β = 1) and *Beta*(*α*=*α_0j_,β*= 1) to model the distribution of *π_i,j_*s when TF *i* regulates gene *j* and when it does not, respectively. However, in order to differentiate between the aforementioned scenarios, we need to impose a restriction on the parameters of these two distributions relative to each other. We hypothesized that the bias towards small values is larger when TF *i* is a regulator of gene *j* compared to when it is not. Intuitively, this can be justified as follows: assuming a linear relationship between the expression of a gene and its regulators, the two main reasons for existence of false positive candidate TFs identified using Elastic Net are the high dimensionality of the data (more features compared to samples), existence of noise in the data and a lack of prior knowledge on the number of regulators of each gene. As a result, even when some false positives are identified using Elastic Net, most of the variance of the gene’s expression is expected to be explained using the expression of the true positive TFs. As a result, the expression of the true positive TFs will have a more significant association with the gene’s expression in an OLS model. We used extensive simulation analysis under different setups and confirmed the intuition above (Supplementary Table S5 and Supplementary Figures S10-S12 in Supplementary File S2). As a result, we modeled the prior distribution of these unknown parameters according to *α*_0*j*_ ~ Unif (0.5,1) and *α*_0*j*_ ~ Unif (0, 0.5), to ensure that *α*_0*j*_>α_1*j*_ and a more significant bias towards small values exists when TF *i* is a regulator of gene *j* (see Supplementary Figure S8). To model the conditional distribution of π_*i,j*_ given its parents, we note that one implication of *T_i,j_* = 1 is that TF *i* regulates gene *j.* On the other hand, if *T_i,j_* = 0, either TF *i* does not regulate gene*j* or TF *i* regulates gene *j* but gene *j* is not associated with the phenotype. Consequently, we used the following model

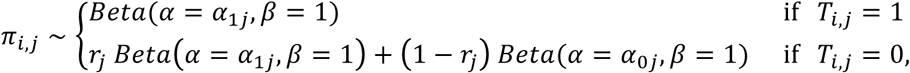

where *r_j_* is an unknown mixing parameter representing the probability that TF *i* regulates gene *j* but gene *j* is not associated with the phenotype. We assigned a prior distribution of *r_j_* ~ *Unif*(0,1) to this parameter (reflecting lack of prior knowledge).

We used a Markov chain Monte Carlo (MCMC) method using the PyMC python module [55] to infer the unknown parameters and learn empirical posterior probabilities for *T_i,j_* s. Since some of the solutions of the MCMC may converge to local optima, to alleviate their effect we ran the MCMC procedure 100 times with different random initializations and obtained an average posterior probability for each *T_i,j_*. These average values were then minmax normalized and an appropriate threshold was used to identify phenotype-relevant regulatory edges (we used a threshold of 0.5). Since several parameters can be configured by the user, for the default values which were used in the pan-cancer analysis as well as the method used for hyperparameter selection see Supplementary Methods (in Supplementary File S2).

### Software availability

Implementations of InPheRNo and simplified-InPheRNo in python, with appropriate documentation, are available at: https://github.com/KnowEnG/InPheRNo and https://github.com/KnowEnG/Simplified-InPheRNo, respectively.

### Data collection and normalization

We downloaded a list of 1544 human TFs from AnimalTFDB [56]. Gene (including TF) expression profiles of 6,357 cancer patients corresponding to 18 different cancer types in TCGA were downloaded from the Genomic Data Commons [22]. Similarly, the gene expression profiles of 4,388 normal tissue samples corresponding to these 18 cancer types (version V6p) were downloaded from the GTEx data portal (www.gtexportal.org). To normalize the FPKM (TCGA) and RPKM (GTEx) values we used an approach similar to the guideline described in the GTEx data portal for analyzing gene expression corresponding to version V6p. The expression profile of each sample was normalized in two ways: for the analyses that involved expression of all samples (across different cancer or tissue types), a pan-cancer (pan-tissue) normalization was performed, while for the analyses that required samples of one cancer (tissue) type, a cancer (tissue) –specific normalization was performed (see Supplementary Methods in Supplementary File S2).

For the comparison of the reconstructed networks using InPheRNo with a global (cancer-agnostic) TRN, we downloaded *“*ENCODE TREG binding profiles*”* from http://eh3.uc.edu/treg which include the binding probabilities assigned to each (TF, gene) by TREG for 43 different cell lines. We then selected edges with probability larger than 0.5 and formed their union over all cell lines to obtain a global TRN.

We obtained from IntOGen (www.intogen.org) a list of driver TFs that are identified based on mutations, gene fusions and copy number alterations. We then combined the driver lists for each of these three data types into one list for each cancer. We also obtained a list of cancer driver genes from http://driverdb.tms.cmu.edu.tw/driverdbv2, selecting driver genes that were identified by at least two different methods.

### Alternative baseline approaches for network reconstruction

We used several alternative approaches to TRN reconstruction as comparators for InPheRNo. The context-restricted network reconstruction generally refers to an approach in which a network reconstruction algorithm is applied to samples of a context of interest, excluding the samples corresponding to other contexts. Since any TRN reconstruction algorithm based on gene expression data can be used in this framework we used Elastic Net [57–59], which we have also used as the first step of InPheRNo, to ensure a fair comparison between InPheRNo and context-restricted network analysis. To obtain a context-restricted network for each cancer type, we used the expression profile of a gene across samples of that cancer type as the response vector and the expression of the TFs as the feature vectors to identify TFs with nonzero coefficients for each gene. Details of choosing the hyperparameters of the Elastic Net using cross-validation are provided in the Supplementary Methods (in Supplementary File S2).

In the differential network analysis, we used the context-restricted analysis described above to reconstruct two networks for each cancer type: one using samples of that cancer and another using samples of all other 17 cancers. Then we constructed a differential network by identifying edges that are present in the former network but not in the latter. In the context-specific approach, we first identified top 1500 genes that were differentially expressed between one cancer type compared to other types of cancer (Bonferroni-corrected p < 1E-20). Then, we used Elastic Net to relate the expression of these genes to the expression of TFs.

In simplified-InPheRNo, we used Pearson’s correlation to obtain p-values of TF-gene correlation and a two-sided t-test to obtain the p-values of gene-phenotype associations differentiating one cancer type from other types of cancer. Next, for each (gene, TF, phenotype) triplet, we used Fisher’s method to combine the two p-values. Then for each cancer type, edges with smallest p-values were selected such that the number of edges in the reconstructed network would be equal to the number of edges identified by InPheRNo (for a fair comparison). We performed this analysis for each cancer type using TCGA data and each tissue type using GTEx data and used the same approach in InPheRNo to remove the confounding effect of tissues of origin.

To ensure the fairness of comparisons, in all except for context-specific analysis, we focused on the same subset of genes that were utilized in analyzing InPheRNo. For context-specific TRNs, we used differential expression analysis to identify putative target genes, since this is part of the method itself.

### Forming gene expression signatures using reconstructed networks for survival analysis

We defined the gene expression signature of a TF in each cancer type as a weighted linear combination (***x***+**∑**_*i*_w_i_**y**_*i*_) of the expression profile of the TF (denoted by *x)* and its targets (denoted by *y_i_*) across different samples of that cancer type. To consider the strength and mode of regulation for each gene, we used the Pearson’s correlation coefficient between the expression profile of the TF and each target gene as the weights (*w_i_* s) in this linear combination. This signature reflects the expression of the TF as well as the activity level of its targets, while considering the mode and strength of regulation. For each TRN reconstruction method, agglomerative clustering (with average linkage) was applied to the signatures of 5 expressed TFs with the most identified targets to cluster samples into two distinct groups for survival analysis. We used cosine similarity with this clustering method since it has been shown to be one of the best options for clustering of cancer samples using gene expression data [60].

## Authors’ Contributions

AE and SS conceived the study and designed the algorithm. AE implemented the algorithm and performed the statistical analyses of the results. AE and SS contributed to the drafting of the manuscript and critical discussion of the results. Both authors read and approved the final manuscript.

## Funding sources

This work was supported by research grant 1U54GM114838 awarded by NIGMS through funds provided by the trans-NIH Big Data to Knowledge (BD2K) initiative. We thank Roy Campbell for providing compute resources for the analyses.

